# Tumor-associated mammary fibroblast matrices contain O-glycosylated oncofetal fibronectin

**DOI:** 10.64898/2026.09.29.755525

**Authors:** Jessica H Longstreth, Deepp Patel, Ghazal Bashiri, Karin Wang

**Author notes:** Corresponding author: 1964 N. 12th Street, Philadelphia, PA 19122 USA.

## Abstract

Oncofetal fibronectin (oFN) is a known biomarker of cancer progression and is involved in events such as epithelial mesenchymal transition and multidrug resistance. It is also understood that fibronectin (FN) and its splice variants are differentially arranged by cancer associated fibroblasts (CAFs) to support cancer cell proliferation and migration. Contrastingly, little is understood about the mechanobiology of oFN, a post-translationally modified FN isoform defined by O-glycosylation of a specific threonine residue. To investigate the architectural features of the oFN matrix, CAFs were simulated by culturing human mammary fibroblasts (HMFs) on stiff polyacrylamide gels (20 kPa) in media collected from HCC-1937 (primary site breast adenocarcinoma) and MDA-MB-231 (metastatic breast adenocarcinoma) cells. As a control, HMFs were also cultured on softer gels representing healthy adipose tissue (2 kPa) in media not exposed to cancer cells. After immunocytochemical analysis, tumor conditioned HMFs displayed an expected CAF-like phenotype that was most prominent with MDA-MB-231 conditioned media. Matrices produced by tumor-conditioned HMFs contained more oFN with structural features mostly following that of the overall dense, aligned FN matrix. Cancer cells reseeded on their corresponding decellularized tumor-associated matrices had greater F-actin/oFN colocalization than control matrix. The lack of distinct oFN topographical cues may indicate that oFN O-glycosylation is more directly involved in biochemical signaling than mechanosignaling through matrix architecture. Together, this study established a platform for investigating stromal oFN which revealed that tumor-conditioned media sourced from cancer cells with different metastatic potential influences HMF activation, leading to distinct total FN and thus oFN matrix microarchitecture features.

**New and Noteworthy:** This study is the first to investigate topographical features of the oFN matrix, utilizing decellularized ECMs to specifically visualize oFN matrix deposition. Unlike other FN variants, Unlike the EDA and EDB splice variants, oFN microarchitecture does not seem to significantly differ from the total FN matrix. This suggests that the O-glycosylation site defining oFN is not intended for direct mechanosignaling, warranting further mechanistic studies of oFN.

## Introduction

Despite advances in treating even aggressive forms of breast cancer (1), it remains the most diagnosed cancer worldwide (2) and multidrug resistance (MDR) in metastatic tumors remains a major challenge (1). While much can be learned from cancer cells themselves, the surrounding tumor microenvironment (TME), made up of the extracellular matrix (ECM) and cancer-associated cells, cannot be ignored. It is well understood that the TME modulates breast cancer progression and recent studies have even suggested its prognostic value (3–5). Within the TME, cancer-associated fibroblasts (CAFs) are generally defined by an excess deposition of ECM fibers, including fibronectin (FN). FN is a key glycoprotein in the ECM, mediating important events in cell pathophysiology (6, 7). Several variants of fibronectin exist and are known to play their own distinct roles in tumor progression (8). Among these variants is oncofetal fibronectin (oFN), which is an O-glycosylated form of FN (9). Although oFN has been implicated in breast cancer MDR, mechanistic studies surrounding oFN are limited.

oFN is found in fetal and adult cancer tissues, but not in healthy adult tissues (10). This variant was first recognized by Hidemitsu Matsuura and Sen-itiroh Hakomori in 1985 after they synthesized monoclonal antibody (mAb) FDC-6 by immunizing mice with FN isolated from HuH-6 hepatoma cells. Interestingly, this antibody was reactive for FNs from placenta, amniotic fluid, fetal tissues, and tumor tissues but did not bind to FNs from various adult tissues. Thus, FN recognized by the FDC-6 antibody was termed “oncofetal” FN (10). Subsequent studies found that this oFN antibody binds to an O-glycan at a specific threonine residue in the type III connecting segment (or variable region) of FN (9). Unlike variants that arise from alternative splicing, oFN is a post-translationally modified FN isoform; normal FN becomes oFN when a glycosyltransferase adds a GalNAc unit to the glycosylation site that defines oFN -- a process called O-glycosylation. It should be noted that extra domain A (EDA) and extra domain B (EDB) splice variants of FN are sometimes referred to as “oncofetal fibronectin” in literature, as these variants are also associated with fetal development and cancer processes (11–13). oFN as discussed here refers to the O-glycosylated form of FN as recognized by the FDC-6 mAb; it can also contain EDA and/or EDB.

In the context of oncology, oFN is a predictive biomarker and is involved in protumorigenic processes. For instance, epithelial mesenchymal transition (EMT) is a well-established hallmark event in cancer progression and metastasis (14). Investigation into oFN’s role in this process revealed that oFN and glycotransferase expression are elevated in TGF-β-induced EMT. Selective siRNA knockdown of the glycotransferases GalNAc-T3 and GalNAc-T6 inhibited TGF-β-induced oFN up-regulation. Mesenchymal markers were also reduced after glycotransferase knockdown, suggesting that O-glycosylation of FN may facilitate EMT (15). Additionally, oFN is upregulated in the MDR phenotype of MCF-7 human breast adenocarcinoma cells. Silencing of pp-GalNAc-T6, the glycotransferase suspected to be mainly responsible for oFN production, resulted in reduced oFN expression and partial reversal of the MDR phenotype (16). oFN also has a long history as a cancer biomarker, with many of the studies prior to de Lima *et al*’s 2011 study (15) focusing on simply identifying the presence of oFN in specific cancer tissues (16–19). In the breast, FDC-6 reactive oFN is not detected in normal tissue but is present in most invasive breast carcinomas (20). More recently, oFN was recognized as a cancer stage-specific biomarker, as its expression increases with stage progression and it is only present in stages 2-4 in hepatocellular carcinoma patients. This also implies oFN may have specific functions associated with protumorigenic events at these stages (21). oFN may have therapeutic utility as well since CAR T cells targeting oFN were able to overcome tumor resistance (22). Regardless of years of studies clearly suggesting oFN participates in cancer progression, the mechanisms by which oFN interfaces with cancer cells remain unclear and it has not been analyzed through a mechanobiology lens.

Consideration of oFN mechanobiology is valuable because it is well known that ECM organization and mechanotransduction pathways can affect gene expression and cell behavior (23). Mechanotransduction refers to the conversion of mechanical stimuli to biochemical signals that trigger signaling pathways in the cell to respond accordingly. Integrins are heterodimeric transmembrane receptors with high affinity for ECM proteins and they act as mediators for mechanotransduction (24, 25). In the TME, CAFs deposit an excess of ECM proteins, contributing to a stiffer environment that cells sense through integrins. This stiff, dense ECM network modulates cancer cell signaling and behavior, which can lead to more persistent cancer cell migration and cell spreading, among other events (26–29). CAFs can also remodel FN through integrin binding, creating more aligned matrices that promote directional migration of cancer cells (30, 31). Queries into other FN variants reveal that EDA and EDB FN also promote oncogenic cell responses and have distinct matrix organization (32–34). These variants also have high affinity to integrins beyond the major FN integrin, α_5_β_1_, and these interactions affect downstream processes such as cell adhesion and differentiation (35, 36). It follows that oFN may also be differentially remodeled by CAFs and have specific integrin interactions that contribute to cancer progression.

An effective technique for investigating TME matrix mechanobiology is the use of decellularized ECMs (dECMS). Sophisticated models of the TME that recapitulate stromal components are necessary since 2D cell culture on plastic substrates causes cells to behave differently than they would *in vivo.* For *in vitro* studies, dECMs offer a biomimetic substrate that provides more relevant components of the TME than synthetic alternatives (37). Many dECMs are derived from whole tissue samples (38–41) and while these matrices provide complex microarchitecture and composition, it can be difficult to isolate specific variables in these complicated model systems (41). Cell-derived dECMs can strike a balance between offering mimetic matrix components and allowing modulation of specific variables in an *in vitro* model. The Cukierman group has successfully used patient derived CAFs to obtain and characterize tumor-associated dECMs (42). CAFs have also been modeled by growing fibroblasts in tumor conditioned media (TCM) and their matrices revealed differences in fibronectin and collagen matrix deposition between tumor conditioned fibroblasts and control group fibroblasts (43, 47). These studies demonstrate the utility of tumor-conditioned media and dECMs for studying matrix microarchitecture that may contribute to cancer progression.

This study aimed to simulate bulk stiffness and tumor secreted factors of the TME in human mammary fibroblast (HMF) culture to produce tumor-associated dECMs (**Fig. 1**). Two different breast adenocarcinoma cell lines were used to produce TCM: HCC-1937 and MDA-MB-231 cells. HCC-1937 cells are from a primary tumor site while MDA-MB-231 cells are from a metastatic pleural effusion, allowing for investigation of how factors secreted by breast cancer cells of different tumor sites and aggressiveness influence HMF activation and FN deposition. HMFs were also grown in conditions more similar to healthy adipose tissue as a control. Immunocytochemical analysis was used to compare total FN and oFN matrix features including matrix area, fiber width, alignment, and colocalization. Cancer cells were also reseeded onto their corresponding dECMs and control dECMs to evaluate the effect on cancer cell morphology. Overall, these experiments were expected to provide insight into distinct mechanobiological features of oFN in tumor-associated matrices that may contribute to protumorigenic cross-talk with cancer cells.

**Figure 1:**
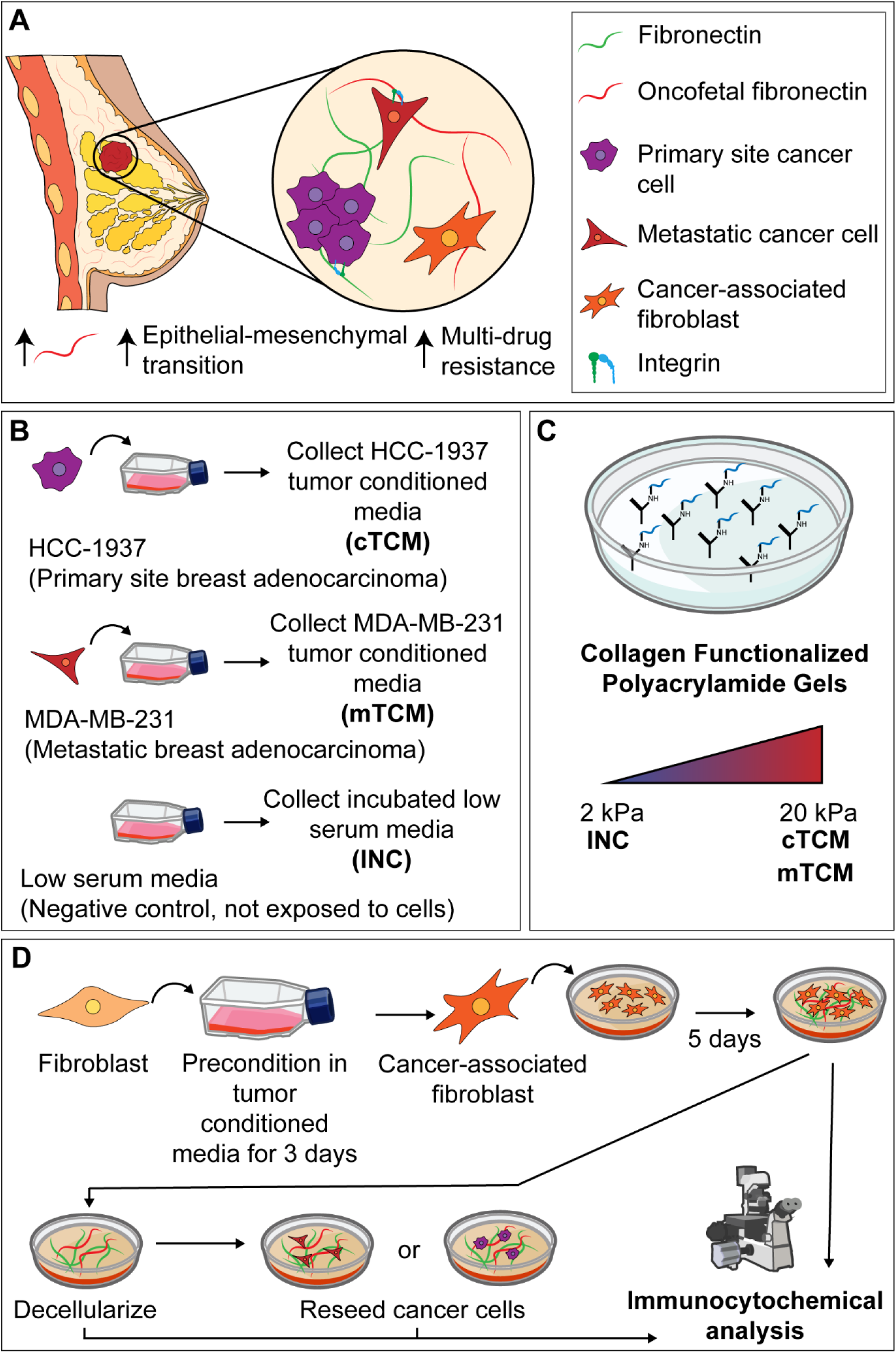
Experimental context and setup. **A)** The breast cancer tumor microenvironment contains a heterogeneous population of cells, including primary site and migrating metastatic cells that release soluble cytokine factors into the environment. Fibroblasts respond to environmental signals and become cancer-associated fibroblasts (CAFs), depositing an excess of ECM proteins, such as oncofetal fibronectin, which contributes to cancerous conditions. Oncofetal fibronectin is associated with epithelial-mesenchymal transition and multi-drug resistance in cancer cells. ECM-integrin binding can activate downstream signaling pathways that affect cancer cell phenotype; however, the mechanisms by which oncofetal fibronectin promotes cancer progression are not well understood. **B)** To recapitulate tumor soluble cues, media exposed to confluent HCC-1937 and MDA-MB-231 cells was collected. **C)** To provide a substrate of relevant bulk stiffness, polyacrylamide gels of healthy (2 kPa) and cancerous (20 kPa) adipose tissue were cast and functionalized with collagen. **D)** Human mammary fibroblasts (HMFs) were grown in tumor conditioned media and seeded on gels of relevant stiffness to model the tumor microenvironment that would promote a CAF phenotype. After 5 days of matrix deposition, samples were either assessed for HMF activation, decellularized for analysis of matrix microarchitecture, or decellularized and reseeded with cancer cells to evaluate the effect of tumor-associated matrices on cancer cell morphology.

## Materials and Methods

### Cell culture

HMFs (ScienCell, 7630), passages 7-10, were used to produce cell-derived matrices in the following experiments. Additionally, breast adenocarcinoma cell lines HCC-1937 (ATCC, CRL-2336, primary ductal carcinoma) and MDA-MB-231 (ATCC, HTB-26, metastatic pleural effusion) were used to produce tumor-conditioned media (TCM) and passage 9-16 were used for reseeding experiments. All cells were cultured in Dulbecco’s Modified Eagle Medium (DMEM) (Gibco, 11995073) supplemented with 10% fetal bovine serum (FBS) (GeminiBio, S11150) and 1% penicillin-streptomycin (Gibco, 15140122).

### Tumor-conditioned media collection

To produce growth media containing relevant soluble factors to breast adenocarcinoma cell lines of varying metastatic potential, HCC-1937 and MDA-MB-231 cells were cultured to 80-85% percent confluency in high serum media (DMEM supplemented with 10% FBS and 1% penicillin-streptomycin). Then, 24 hours before collection of TCM, cells were given low serum media (DMEM supplemented with only 1% FBS and 1% penicillin-streptomycin) to encourage release of tumor-derived soluble factors versus cell growth. On the day of collection, cells were trypsinized (Gibco, 25200056) and counted using a hemocytometer. Collected media was diluted with fresh low serum media to a concentration of 15 mL of media per 7 million cancer cells based on the cell count. TCM was concentrated 10x using 3 kDa molecular weight cut-off centrifugal filter tubes (Millipore, UFC9003) at 4000 rpm, 4 °C and then frozen at -20 °C for long term storage. TCM from HCC-1937 cells is referred to as “cTCM” while TCM from MDA-MB-231 cells is referred to as “mTCM”.

To produce control group cell culture media, low serum media was incubated for 24 hours at 37 °C, then concentrated to 10x using centrifugal filter tubes at 4000 rpm, 4 °C and frozen at -20 °C for long term storage. This incubated low serum media is referred to as “INC”. All collected media was thawed, diluted to 1.5x in additional low serum media, and sterile filtered through a 0.2 μm filter prior to use in cell culture.

### TGFβ1 enzyme-linked immunosorbent assay

The concentration of TGFβ1 in INC, cTCM, and mTCM was quantified via an enzyme-linked immunosorbent assay (ELISA) kit (R&D Systems, DY240-05), following manufacturer instructions. This kit used a sandwich ELISA approach by which a 96-well plate was coated overnight with capture antibody specific to TGFβ1. Latent TGFβ1 in media samples was activated by acidification then neutralized. Standard solutions of known TGFβ1 concentrations and media samples (3 technical replicates of 3 separate batches of each media type) were loaded into the wells and after an incubation period, captured TGFβ1 was sandwiched with detection antibody. Streptavidin-HRP was added and subsequent addition of TMB substrate followed by acidic stop solution provided a concentration-dependent color change. Absorbance was measured using a TECAN Infinite 200 Pro microplate reader at 450 nm with a wavelength correction reading at 540 nm. The absorbance data was used to make a standard curve from which TGFβ1 concentration in the media samples was determined.

### Fibroblast-mediated matrix assembly

To provide fibroblasts with a substrate of pathophysiological relevant bulk stiffness, polyacrylamide gels were prepared (44, 45) with stiffnesses of 2 kPa and 20 kPa, representing soft, healthy adipose tissue and stiffened, cancerous adipose tissue, respectively. First, the 20 mm glass bottoms of 35 mm petri dishes (Cellvis, D35-20-1.5-N) were given a 20 min acid wash in 2 N H_2_SO_4_, washed three times in DI water, followed by a 20 min incubation with 0.1 M NaOH and an additional three washes of DI water. After drying the dishes with compressed air, the insets were treated with a silane binding solution of 20% 3-(Trimethoxysilyl) propyl methacrylate (Sigma-Aldrich, 440159-100M) and 30% glacial acetic acid (FisherScientific, AA33252AK) in DI water for 1 hour at room temperature. Dishes were then washed 3 times with pure ethanol, another 3 washes with DI water, and then dried again. Gel precursor solutions were made according to final acrylamide (Bio-Rad, 1610140) and bis-acrylamide (Bio-Rad, 1610142) ratios previously defined and verified via atomic force microscopy by Young *et al* (46) to achieve the 2 kPa and 20 kPa stiffnesses. Ammonium persulfate (10% w/v, Bio-Rad, 1610700) and N,N,N’,N’-tetramethylethylenediamine (Bio-Rad, 1610800) crosslinkers were added to precursor solutions at a 1:100 and 1:1000 ratio to initiate gel formation. 24 uL of gel solution were added to each inset and sandwiched under an 18 mm circular glass coverslip pretreated with Sigmacote (MilliporeSigma, SL2-100ML) siliconizing agent according to manufacturer instructions. 2 kPa gels and 20 kPa gels were cured in a desiccator for 15 and 20 minutes, respectively before covering in 2 mL of DI water for storage no longer than 3 weeks at 4 °C.

Gels were prepared for collagen functionalization by removing coverslips and adding 400 μL of 0.5 mg/mL sulfo-SANPAH (ThermoFisher, 22589) in 50 mM HEPES (MilliporeSigma, H3375-25G) pH 8.5. The dishes were then exposed to UV light for 15 minutes to activate sulfo-SANPAH and sterilize the gels. During this process, 3 mg/mL bovine collagen (Advanced BioMatrix, 5005-100ML) was diluted to 0.2 mg/mL in sterile PBS pH 7.4 (Corning, 21-040-CV). After activation, excess sulfo-SANPAH was gently removed via three washes of sterile-filtered 50 mM HEPES pH 8.5. 700 μL of 0.2 mg/mL collagen solution was added to each gel and incubated overnight at 4 °C. The following day, collagen treated gels were washed three times with sterile PBS and stored in additional PBS at 4 °C. Functionalized gels were used within three days for fibroblast-mediated matrix assembly.

Three days prior to seeding fibroblasts onto collagen functionalized gels, HMFs were conditioned in appropriate media type (INC, cTCM, or mTCM) (43, 47) to promote phenotypic changes before matrix assembly on the gels. Then, HMFs were trypsinized and 300,000 cells in 0.4 mL of low serum media were seeded into each glass inset. INC-conditioned fibroblasts were seeded on 2 kPa gels while cTCM- and mTCM-conditioned fibroblasts were seeded onto 20 kPa gels. Cells were allowed to settle onto the gels for 20 minutes at room temperature, and then carefully placed into the 37 °C incubator to continue forming nascent adhesions. Once cells were visibly attached to the gels, additional media of the appropriate condition (INC, cTCM, or mTCM) was added to a volume of 2 mL. After 24 hours of additional incubation, media was removed and replaced with the appropriate media type supplemented with 75 μg/mL L-ascorbic acid (FisherScientific, AAJ61899AK) in dPBS (Gibco, 14190250) to encourage matrix deposition. On day 3 of culture, only half of the media was removed and replaced with 150 μg/mL L-ascorbic acid in dPBS (to bring to a final concentration of 75 ug/mL in the dish) (48, 49). On day 5, samples were either fixed for 1 hour in 4% paraformaldehyde in PBS for later immunocytochemical analysis of HMF phenotype or underwent matrix decellularization.

### Matrix decellularization

Extraction buffer was previously made by sterile filtering a solution of 0.5% Triton-X 100 (FisherScientific, MTX15681) and 20 mM NaOH in dPBS (48). After 5 days of HMF-mediated matrix assembly, samples were gently washed three times with dPBS and treated with 1 mL of extraction buffer prewarmed at 37°C. Samples were incubated at 37 °C for 5 minutes and cell lysis was observed under a light microscope. An additional 2 mL of dPBS were slowly pipetted along the side of the petri dish, being careful not to disturb the fragile matrices. The freshly decellularized matrix samples were kept at 4 °C overnight to allow stabilization of the matrices before subsequent DNase treatment and debris washes.

The next day, decellularized matrices were treated with DNase I. Prior to treatment, 10x DNase reaction buffer was made, consisting of 100 mM Tris HCl pH 7.5, 25 mM MgCl_2_, and 1 mM CaCl_2_ in DI water. 10x DNAse inactivation buffer, consisting of 100 mM Tris-HCl in DI water, was also made. All buffers were sterile filtered through 0.2 μm membrane. On the day of DNase treatment, buffers were diluted to 1x in autoclaved sterile filtered DI water and prewarmed at 37 °C. The liquid volume was carefully removed from samples previously stored at 4 °C. One wash of dPBS was slowly added to the side of the petri dish and then removed to reduce cellular debris. DNase I (Invitrogen, 18047019) was added to prewarmed DNase reaction buffer to a final concentration of 10 U/mL and 1 mL was added to each dish. Matrices were incubated with DNase treatment for 30 minutes at 37 °C. Then, the DNase reaction buffer was carefully aspirated from each sample and a 15 minute room temperature incubation with 1 mL DNase inactivation buffer was performed, followed by another two washes with DNase inactivation buffer.

At this point, samples were either fixed for 1 hour in 4% paraformaldehyde in PBS for later immunocytochemical analysis of matrix microarchitecture or stored in dPBS with 1% penicillin-streptomycin at 4 °C for future reseeding experiments (within 3 weeks).

### Cancer cell reseeding on decellularized fibroblast-derived matrices

HCC-1937 and MDA-MB-231 cells were grown to 80-90% confluency before trypsinization and reseeding onto previously prepared decellularized matrices. HCC-1937 cells were seeded onto parallel INC and cTCM matrices, while MDA-MB-231 cells were seeded onto parallel INC and mTCM matrices. All cells were seeded at a density of 20,000 cells, initially seeding cells in 0.1 mL of low serum media pipetted onto the edge of the matrix covered gel and gently swirled to cover the inset. Cells were allowed to settle for 20 minutes at room temperature, then incubated for 30 minutes at 37 °C. Once cells were visibly attached, additional low serum media was added to bring the volume to 2 mL. Samples were incubated for 24 hours at 37 °C, then washed three times with dPBS and fixed for 1 hour in 4% paraformaldehyde in PBS.

### Immunocytochemistry

All samples were washed two times with 0.05% Triton-X 100 in dPBS with gentle agitation for 5 minutes to permeabilize cell membranes. Then, samples were blocked for 30 minutes at room temperature with gentle agitation using 1% SuperBlock (FisherScientific, 37515) and 0.05% Triton-X 100 in dPBS to prepare for antibody treatment. Primary antibodies were incubated overnight at 4 °C. The next day, two 5 minute washes with gentle agitation of 1% SuperBlock and 0.05% Triton-X 100 in dPBS were performed. Fluorophore-containing secondary antibodies and specific dyes were added for a 1 hour room temperature incubation with gentle agitation. Three more 5 minute washes of 1% SuperBlock and 0.05% Triton-X 100 in dPBS with gentle agitation finalized the immunostaining process. Samples were covered with dPBS and sealed with parafilm for long term storage at 4 °C.

For samples with fibroblasts still intact, cell nuclei, F-actin, αSMA, and total FN matrix were labeled. Primary antibodies for αSMA (Invitrogen, 149760-82) and FN (Sigma-Aldrich, F3648) were used at 2 μg/mL and 1:400, respectively. Secondary antibodies, goat anti-mouse Alexa Fluor 647 (Invitrogen, A32728) and goat anti-rabbit Alexa Fluor 488 (Invitrogen, A32731), were used at a 1:100 ratio. Cell nuclei were stained with DAPI (1:5000; Invitrogen, D1306) and F-actin was stained with Alexa Fluor 568 phalloidin (1:250; Invitrogen, A12380).

For decellularized samples, DAPI staining was also performed as described above to verify the removal of cells. The total FN matrix was labeled using the antibody scheme described above. oFN was tagged using 1 ug/mL monoclonal antibody (Creative Biolabs, AGM-190YJ) and fluorescently labeled using secondary antibody goat anti-mouse Alexa Fluor 647.

For samples with reseeded cancer cells, cell nuclei, F-actin and oFN were stained as previously described. All samples were imaged using Olympus Fluoview FV1200 confocal microscope with 30x oil objective. Images of four regions of interest per inset were captured.

### Image analysis

For analysis of HMF phenotype in samples with intact fibroblasts, maximum projection images of cell nuclei, F-actin, αSMA, and total FN were analyzed. Cell nuclei were segmented using Cellpose version 3.1.1.2 and .zip files of segmented nuclei ROIs were exported to Fiji/ImageJ for measurement of cell nuclei area via the ROI Manager feature (50, 51). A custom Fiji/ImageJ macro was written to batch process fluorescent signal area coverage of respective F-actin, αSMA, and total FN channels. Briefly, Subtract Background with a 20 pixel radius was applied, then the image was converted to a 16 bit binary image, converted to an RGB image, and then the Color Pixel Counter plugin was used to identify the number of pixels containing signal in the image and later converted to μm^2^ (52). FN fiber alignment in each image was found using CTFire, which uses a curvelet transform analysis to extract individual fibers from the image, and then CurveAlign, which reports their alignment (53, 54).

For decellularized matrix analysis, fluorescent signal coverage of the total FN and oFN channels was determined using the same Fiji/ImageJ macro described above. Width analysis of total FN and oFN fibers was reported from CTFire results and alignment was reported from CurveAlign results. Colocalization analysis of the binary images of the total FN and oFN channels was achieved using BIOP JACoP ImageJ plugin and the M1 Mander’s Coefficient was reported (55), which represents the fraction of oFN signal overlapping with the total FN matrix.

Cell morphology of cancer cells reseeded onto decellularized matrices was evaluated using Cellpose to segment cell bodies from 2-channel images of the cell nuclei and F-actin channels. These were exported to the ImageJ ROI manager for measurement of single cell area and circularity. Colocalization of the binarized F-actin and oFN channels of each image was determined using the BIOP JACoP ImageJ plugin and the Mander’s M1 coefficient was reported, representing the fraction of F-actin signal overlapping with oFN signal.

### Statistical analysis

At least three samples and four regions of interest per sample were analyzed for all experiments. For all tests, α = 0.05. Separate Brown-Forsythe one-way ANOVA tests with post-hoc Dunnett T3 tests were used to compare average cell nuclei area, αSMA fluorescent signal coverage, F-actin fluorescent signal coverage, FN fluorescent signal coverage, and FN fiber alignment between INC, cTCM, and mTCM groups. TGFβ1 concentration in each media type was compared via one-way ANOVA. Separate two-way ANOVA tests with post-hoc Tukey test were used to compare total FN and oFN matrix features (fluorescent signal coverage, alignment, and fiber width) between groups. Colocalization of the oFN and total FN matrix signals were evaluated via Brown-Forsythe one-way ANOVA test with post hoc Dunnett T3 test. Single-cell area and circularity of cells on tumor-conditioned matrices versus INC matrices were evaluated via Welch’s t-test. Colocalization of f-Actin and oFN matrix signals from cells reseeded on tumor conditioned matrices versus INC matrices were evaluated via Welch’s t-test. All statistical analyses were performed using GraphPad Prism software.

## Results

### Protumorigenic cues enhance human mammary fibroblast activation

HMFs were cultured on polyacrylamide gels of increased pathological stiffness (20 kPa) and cultured in TCM to assess the effect of combined protumorigenic cues on HMF phenotype (**Fig. 2A**). TCM from two different breast adenocarcinoma cell lines was used: cTCM from HCC-1937 cells (originally from a primary site tumor) and mTCM from MDA-MB-231 cells (a metastatic line from a pleural effusion). As a control, additional HMFs were cultured on gels with a bulk stiffness representative of healthy adipose tissue (2 kPa) and cultured in INC media, a low-serum media not exposed to cancer cells. After immunocytochemical analysis, images of cTCM and mTCM groups had significantly more αSMA fluorescent area coverage than the INC group (**Fig. 2C**). Average cell nuclei area and F-actin fluorescent area were greater in HMFs in mTCM conditions than in cTCM and INC conditions (**Fig. 2B, 2D**), suggesting more drastic cytoskeletal changes. This trend was mirrored in the FN deposition of mTCM-HMFs; FN fluorescent signal coverage and alignment were elevated compared to other groups (**Fig. 2E-F**). Interestingly, this aligns with the trend in TGFβ1 concentration across INC, cTCM, and mTCM media types. ELISA results indicated a greater TGFβ1 content in mTCM, although not by a significant amount with the sample size tested (**Fig. 2G**). Together, these findings imply that protumorigenic cues modeled by the *in vitro* experimental system produce HMF activation characteristic of cancer-associated fibroblasts (56).

**Figure 2:**
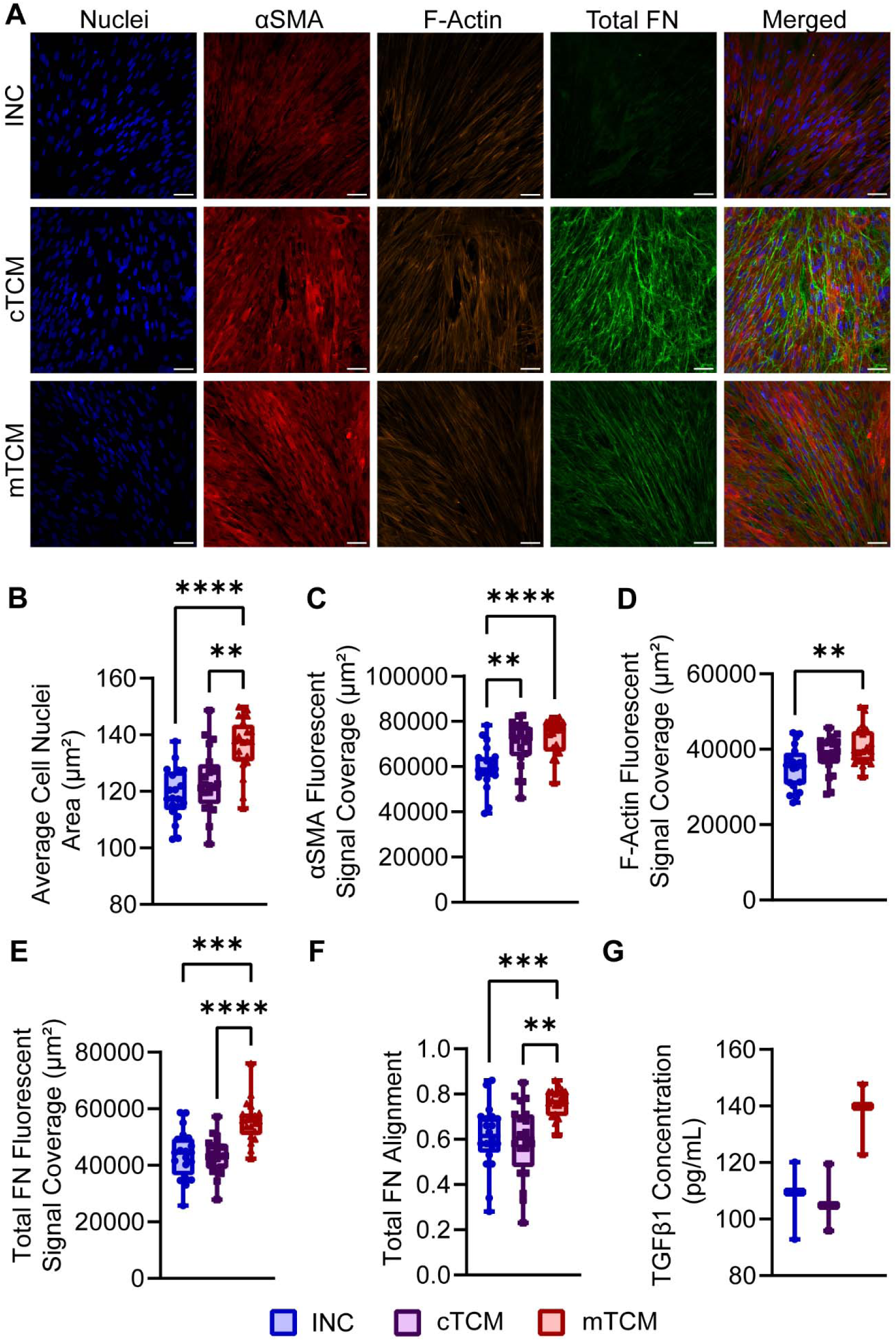
Tumor-conditioned human mammary fibroblasts exhibit a cancer-associated fibroblast-like phenotype. **A)** Representative maximum projection images of human mammary fibroblast (HMF) cell nuclei (blue), alpha smooth muscle actin (αSMA) (red), F-Actin (orange), total fibronectin (FN) (green) and the merged image of all channels for INC, cTCM, and mTCM groups. INC refers to HMFs cultured on 2 kPa polyacrylamide gels and fed with incubated low serum media not exposed to cancer cells. cTCM and mTCM HMFs were cultured on 20 kPa gels fed with tumor-conditioned media collected from HCC-1937 and MDA-MB-231 breast adenocarcinoma cells, respectively. Scale bars 50 μm. **B)** Average cell nuclei area of groups. Cell nuclei for each image were segmented using Cellpose and exported outlines were measured in ImageJ’s ROI manager to find the average cell nuclei area for the region of interest. Four regions of interest from five independent replicates were analyzed. **C-E)** αSMA, F-actin, and FN fluorescent signal coverage, respectively. Fluorescent signal area was determined by counting color-containing pixels in binarized maximum projection images of each channel. Each data point represents a region of interest from five independent replicates. **F)** Total FN alignment, quantified by CurveAlign after using CTFire’s curvelet-transform based extraction of fibers. Each data point represents a region of interest from five independent replicates. **G)** TGFβ1 concentration for each media type. Each data point is the average of three technical replicates, with three independent experiments tested for each media type. For all plots, error bars represent interquartile range and middle line represents median. Asterisks denote statistical significance after hypothesis testing: * p < 0.05, ** p < 0.01, *** p *< 0.001, **** p < 0.0001*.

### Matrices of tumor-conditioned human mammary fibroblasts are differentially organized

Additional HMFs were cultured in the aforementioned INC, cTCM, and mTCM conditions and after 5 days of matrix deposition, samples were decellularized, leaving the conditioned matrices behind. Samples were immunostained for FN and the oFN isoform to allow image analysis of total FN and oFN matrix microarchitecture. DAPI staining was performed to verify successful decellularization; visually, little nuclear debris remained after cell lysis and DNase treatment (**Fig. 3A**). Tumor-associated matrices (deposited by cTCM and mTCM conditioned HMFs) contained greater amounts of oFN than control INC matrices, as indicated by fluorescent signal coverage (**Fig. 3B**). Similar to FN fiber alignment trends when HMFs were left intact, the total FN fibers of mTCM matrices were much more aligned than fibers of cTCM and INC matrices. oFN fibers generally followed the same alignment trends as the total FN matrix across groups (**Fig. 3C**). Fiber width of total FN fibers decreased with tumor conditioning and oFN fiber width for the mTCM group was lower than the cTCM group (**Fig. 3D**). Colocalization analysis of the oFN and total FN matrix revealed a smaller fraction of the oFN signal overlapped with the total FN matrix in cTCM matrices than in INC matrices (**Fig. 3E**). Analysis of total FN and oFN matrix features illustrate that matrices deposited by tumor-conditioned fibroblasts are architecturally distinct from those deposited by less activated, control fibroblasts.

**Figure 3:**
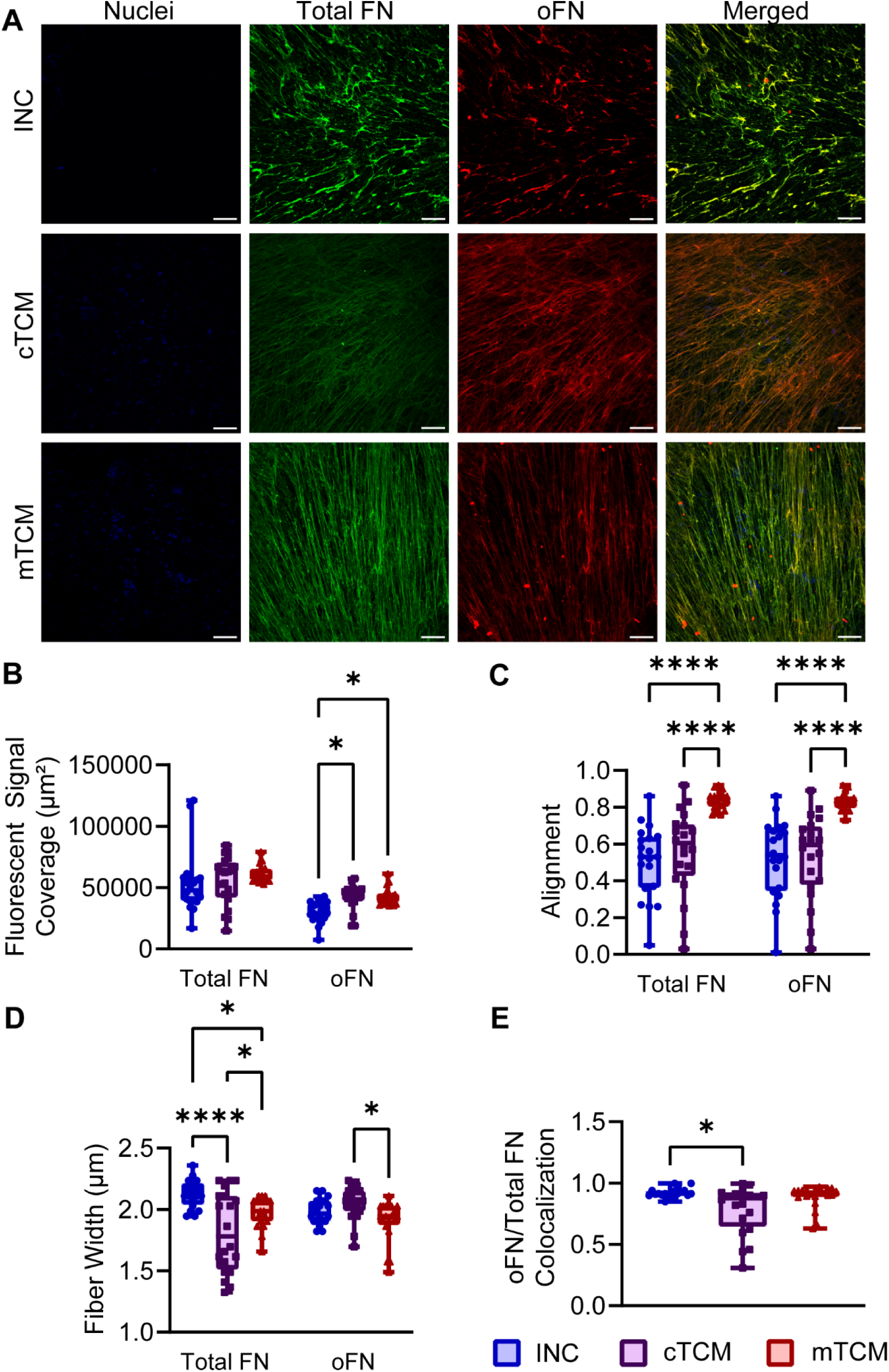
Tumor-conditioned matrices contain oFN with distinct microarchitecture. **A)** Representative maximum projection images of HMF-derived decellularized matrices; cell nuclei debris (blue), total FN (green), oFN (red), and merged image of all channels are displayed. Scale bars 50 μm. **B)** Total FN and oFN fluorescent signal coverage for INC, cTCM, and mTCM groups. Fluorescent signal area was determined by counting color-containing pixels in binarized maximum projection images of each channel. **C)** Total FN and oFN alignment, quantified by CurveAlign after using CTFire’s curvelet-transform based extraction of fibers. **D)** Total FN and oFN fiber width from CTFire analysis. **E)** Colocalization of oFN and total FN, where the Mander’s M1 coefficient represents the fraction of oFN signal overlapping with total FN signal. Colocalization analysis was performed using BIOP JACoP ImageJ plugin. For all plots, each data point represents one region of interest from five independent replicates. Error bars represent interquartile range and middle line represents median. Asterisks denote statistical significance after hypothesis testing: * p < 0.05, ** p < 0.01, *** p < 0.001, **** p < 0.0001.

### Tumor-conditioned matrices influence cancer cell morphology

HCC-1937 and MDA-MB-231 breast adenocarcinoma cells were reseeded onto their corresponding tumor associated matrices and INC control matrices to evaluate the effect of oFN matrix organization on cancer cell morphology. After 24 hours, immunocytochemical analysis was performed and cell bodies were segmented from maximum projection images of regions of interest (**Fig. 4A, 4E**). For HCC-1937 cells, there was no difference in cell area or circularity of cells seeded on cTCM matrices versus INC matrices (**Fig. 4B-C**). The more metastatic MDA-MB-231 cells reseeded on mTCM matrices did have a slightly enlarged cell area compared to MDA-MB-231 cells on INC matrices (**Fig. 4F**), but no difference in circularity (**Fig. 4G**). Overlap between F-actin and oFN signal for each image was also measured via Mander’s M1 coefficient, as it is well documented that FN and actin filaments become more colocalized as cell integrins bind to FN and form focal adhesions (57–59). For both HCC-1937 and MDA-MB-231 cells, there was a greater colocalization of F-actin and oFN for cells reseeded on tumor-associated matrices than INC matrices (**Fig. 4D, 4H**). One outlier was removed from the HCC-1937 colocalization dataset according to ROUT method with a Q value of 1%. These results suggest that cancer cells have greater interactions with oFN in tumor-conditioned matrices. Metastatic potential of the cancer cell may also contribute to cell morphology response.

**Figure 4:**
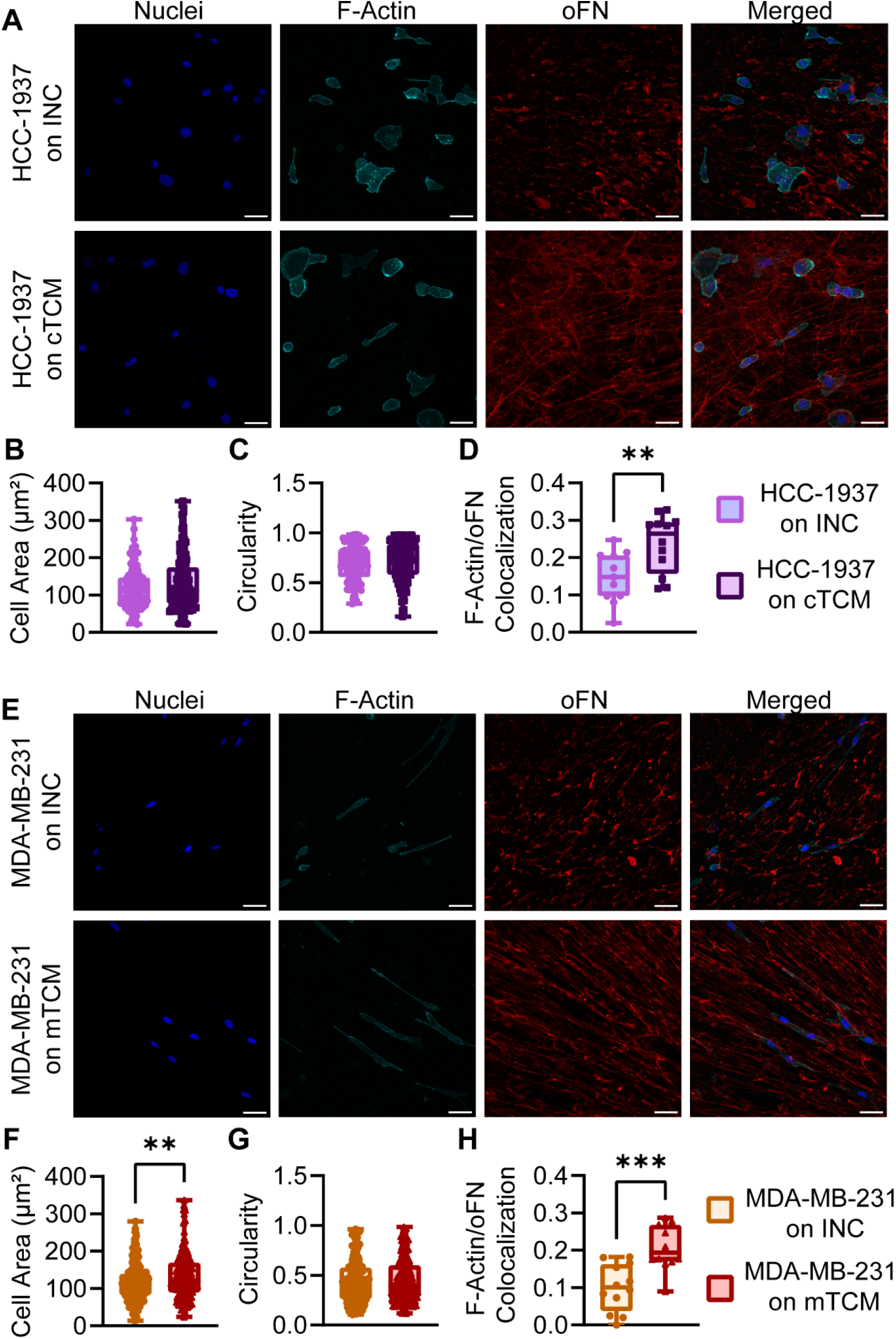
Breast adenocarcinoma cells on tumor-conditioned oFN matrices display different morphologies. **A and E)** Representative maximum projection images of HCC-1937 and MDA-MB-231 cells, respectively, reseeded onto INC and respective tumor-associated matrices; cell nuclei (blue), F-actin (cyan), oFN (red), and merged image of all channels are displayed. Scale bars 50 μm. **B and F)** Cell area of HCC-1937 and MDA-MB-231 cells, respectively. Combined nuclei and F-actin channel images were segmented using Cellpose and area of exported cell outlines were measured in ImageJ. Each data point represents a single cell. **C and G)** Circularity of HCC-1937 and MDA-MB-231 cells, respectively, measured from cell outlines. Each data point represents a single cell. **D and H)** Colocalization of F-actin and oFN signals for HCC-1937 and MDA-MB-231 samples, respectively. Displayed colocalization coefficient is the Mander’s M1 value, which is the fraction of F-actin signal overlapping with oFN signal. Each data point is one region of interest from three independent replicates. For all plots, error bars represent interquartile range and middle line represents median. Asterisks denote statistical significance after hypothesis testing: * p < 0.05, ** p < 0.01, *** p < 0.001, **** p < 0.0001.

## Discussion

This study utilized bulk stiffness and tumor-secreted cytokine cues to mimic the breast cancer tumor microenvironment, creating a conditioned fibroblast-derived matrix platform to better understand oFN matrix characteristics and function. Tumor-conditioned HMFs displayed activation markers characteristic of CAFs, namely increased αSMA compared to the control group. While CAFs have been revealed to consist of heterogenous subpopulations with variable αSMA levels in different contexts (56, 60, 61), αSMA-positive CAFs are generally found across several subpopulations, including matrix and myofibroblast-like CAFs (62, 63). Co-culture of αSMA-positive CAFs increased breast cancer cell growth and overexpression of these CAFs correlates with poor prognosis in cancer patients (64, 65). It should be noted that control group INC HMFs also had visible αSMA signal, but healthy adipose tissue typically has few αSMA-positive cells (66). This is likely due to the limitations of an *in vitro* system; cells are exposed to more activating factors present in FBS and may retain memory of stiff culture flask substrate (67, 68). Despite this, the increased expression of αSMA in HMFs in cTCM and mTCM conditions suggests that the *in vitro* system successfully simulated TME factors, driving tumor-conditioned HMFs further towards a CAF-like phenotype.

mTCM HMFs showed even more evidence of activation, with greater nuclear area and F-actin signal. Nuclear deformation is another hallmark of the transition to a CAF phenotype, as enlargement of nuclear pores due to cytoskeletal forces allows yes associated protein (YAP) to enter the nucleus and upregulate CAF-related genes (69, 70). The increased F-actin signal in mTCM conditioned HMFs could be an indication of cytoskeletal strain that, when transferred to the nucleus through LINC complexes, results in nuclear flattening (71, 72) consistent with the larger average nuclear area observed in this study. This mechanotransduction pathway is believed to reinforce itself: the stiff environment is sensed by F-actin networks, leading to downstream YAP translocation, expression of contractile CAF markers, and heightened sensitivity to the stiffened microenvironment (70). The increased fibronectin deposition by the mTCM group also plays into this CAF activation loop, as enhanced deposition of ECM fibers that contribute to overall stiffening of the microenvironment is another characteristic of CAFs (56, 73). CAFs have also been shown to remodel FN matrices into aligned tracks to promote directional cancer cell migration (31) which may explain the increased FN matrix alignment in the mTCM group. The absence of these additional CAF indicators by any significant measure in the cTCM group suggests that metastatic potential of the cancer cell impacts CAF recruitment. Consistent with this is the observation that TGFβ1 concentration in cTCM (collected from HCC-1937 cells) was similar to that of INC media which was not exposed to any cancer cells. As TGFβ1 is a major contributor to myoCAF activation (74, 75), it follows that the cTCM conditioned HMFs (exposed to less TGFβ1) would be less activated than mTCM HMFs. It has also been demonstrated that the TME of metastatic tumors have a higher CAF proportion than primary site tumors and that CAFs can increase the metastatic ability of cancer cells through a variety of mechanisms (76), all of which further supports increased CAF activation in the mTCM group.

Decellularized tumor-associated matrices also exhibited increased expression of oFN, an O-glycosylated isoform of FN that contributes to cancer progression. Interestingly, the oFN matrix features largely followed that of the overall FN matrix, concerning alignment and fiber width. Once again, the mTCM group had greater matrix alignment which correlates with the increased metastatic potential of the MDA-MB-231 cell line. Fibers were also thinner, which could be indicative of increased strain on the matrix due to contractile tumor conditioned HMFs stretching and unfolding the fibronectin domains at early time points before fibers can be reinforced into thicker anisotropic bundles (77, 78). Alternatively, it is possible that the matrix has developed enough that the FN matrix is being remodeled and load is being transferred to the collagen I matrix as it uses the initial FN matrix as a template (43). Further timeline experiments and FRET analysis would need to be performed to confirm these speculations.

oFN was less colocalized with the total FN matrix in the cTCM group. It is not uncommon for FN variants to be differentially remodeled as variant forms can have different structure-function relationships than the overall FN matrix. For instance, EDA and EDB FN have been found to have different topographical matrix features that contribute to differing downstream cellular responses (34). Yet the oFN matrix in the mTCM group more closely resembled the overall FN matrix. This makes some sense, as the matrix was more aligned and so both fiber types would be positioned more similarly. Once again, this could contribute to the directed migration of the more metastatic cell line that the mTCM matrices are associated with (31). But since the overall FN matrix also follows these trends, this does not suggest that the oFN matrix is responsible for specific architectural cues that promote cancer progression. Instead, the lack of distinct oFN structural features in tumor associated matrices suggests that this glycosylated form of fibronectin is perhaps more involved in downstream biochemical signaling that can explain its known protumorgenic effects. This is supported by the increase in colocalization of cancer cell F-actin and oFN matrix on tumor associated matrices, as the colocalization of F-actin and matrix proteins may indicate increased interactions of the cell and oFN matrix at focal adhesion sites (79). While FN splice domains are known to interact with variant-specific integrins (35, 36), it may be more appropriate to consider galectin signaling at the oFN glycosylation site. Galectins are a class of proteins, often located within or on cells, that bind to specific glycans (80). Since oFN is defined by a glycosylation site, it is possible that the function of this FN modification is to modulate galectin signaling. In fact, galectin-3 (Gal3) is also upregulated in the metastatic niche; it has been proposed as a diagnostic marker for thyroid cancer (81) and found to associate with fibronectin, promoting metastasis in tissues with aberrant glycosylation (82, 83). Galectin binding regulates integrin turnover (80). In FN remodeling specifically, Gal3 stimulates α_5_β_1_ activation through the focal adhesion kinase and phosphatidylinositol 3-kinase pathway (84). This presents an unexplored role for oFN in galectin-dependent signaling.

Together, this study validated an *in vitro* platform for investigating oFN deposition in the simulated tumor microenvironment. HMFs in this system displayed expected signs of activation characteristic of CAFs and produced matrices containing oFN. The matrix microarchitecture and response of cancer cells reseeded on these matrices depended heavily on the metastatic potential of the cancer cell line used to produce the tumor associated matrix, with MDA-MB-231 cells showing the greatest morphological changes on dense, aligned FN matrices. It was important to investigate the structure of the oFN matrix as it is becoming increasingly clear that oFN is involved with cancer progression, yet little is known about its mechanobiological role. Analysis of oFN matrix features shows that it largely tracks with the overall FN matrix, which is not necessarily true of extra-domain splice variants of FN. This suggests that the O-glycosylation site on oFN may be involved in more biochemical cell signaling pathways, such as galectin binding, rather than direct mechanical cues. These potential pathways should be studied further to evaluate whether there are worthwhile therapeutic targets for interrupting detrimental oFN signaling.

## Acknowledgements

KW acknowledges startup funds from the Bioengineering Department at Temple University, an Office of the Vice President for Research Bridge grant, an Office of the Vice President for Research Blue Sky Initiative grant, and a grant (#2305) from the W.W. Smith Charitable Trust. JL and GB acknowledge Teaching Assistantships from the Bioengineering Department at Temple University. DP acknowledges the Temple University Engineering Undergraduate Research Program. The authors thank the shared departmental resources at Temple University for equipment usage.

## References

1. Swain SM, Shastry M, Hamilton E. Targeting HER2-positive breast cancer: advances and future directions. Nat Rev Drug Discov 22: 101–126, 2023. doi: 10.1038/s41573-022-00579-0.

2. Arnold M, Morgan E, Rumgay H, Mafra A, Singh D, Laversanne M, Vignat J, Gralow JR, Cardoso F, Siesling S, Soerjomataram I. Current and future burden of breast cancer: Global statistics for 2020 and 2040. Breast 66: 15–23, 2022. doi: 10.1016/j.breast.2022.08.010.

3. Danenberg E, Bardwell H, Zanotelli VRT, Provenzano E, Chin S-F, Rueda OM, Green A, Rakha E, Aparicio S, Ellis IO, Bodenmiller B, Caldas C, Ali HR. Breast tumor microenvironment structures are associated with genomic features and clinical outcome. Nat Genet 54: 660–669, 2022. doi: 10.1038/s41588-022-01041-y.

4. Domínguez-Cejudo MA, Gil-Torralvo A, Cejuela M, Molina-Pinelo S, Salvador Bofill J. Targeting the Tumor Microenvironment in Breast Cancer: Prognostic and Predictive Significance and Therapeutic Opportunities. Int J Mol Sci 24: 16771, 2023. doi: 10.3390/ijms242316771.

5. Tian Y, Yang Y, He L, Yu X, Zhou H, Wang J. Exploring the tumor microenvironment of breast cancer to develop a prognostic model and predict immunotherapy responses. Sci Rep 15: 12569, 2025. doi: 10.1038/s41598-025-97784-9.

6. Hynes R. Molecular Biology of Fibronectin. Ann Rev Cell Biol 1: 67–90, 1985. 10.1146/annurev.cb.01.110185.000435.

7. Longstreth JH, Wang K. The role of fibronectin in mediating cell migration. American Journal of Physiology-Cell Physiology 326: C1212–C1225, 2024. doi: 10.1152/ajpcell.00633.2023.

8. Rick JW, Chandra A, Dalle Ore C, Nguyen AT, Yagnik G, Aghi MK. Fibronectin in malignancy: Cancer-specific alterations, protumoral effects, and therapeutic implications. Semin Oncol 46: 284–290, 2019. doi: 10.1053/j.seminoncol.2019.08.002.

9. Matsuura H, Takio K, Titani K, Greene T, Levery SB, Salyan MEK, Hakomori S. The Oncofetal Structure of Human Fibronectin Defined by Monoclonal Antibody FDC-6: Unique Structural Requirement for the Antigenic Specificity Provided by a Glycosylhexapeptide. Journal of Biological Chemistry 263: 3314–3322, 1988.

10. Matsuura H, Hakomori S-I. The oncofetal domain of fibronectin defined by monoclonal antibody FDC-6: Its presence in fibronectins from fetal and tumor tissues and its absence in those from normal adult tissues and plasma (hepatoma/COOH-terminal region). Proceedings of the National Academy of Sciences 82: 6517–6521, 1985.

11. Kumra H, Reinhardt DP. Fibronectin-targeted drug delivery in cancer. Adv Drug Deliv Rev 97: 101–110, 2016. doi: 10.1016/j.addr.2015.11.014.

12. Hall RC, Vaidya AM, Schiemann WP, Pan Q, Lu Z-R. RNA-Seq Analysis of Extradomain A and Extradomain B Fibronectin as Extracellular Matrix Markers for Cancer. Cells 12: 685, 2023. doi: 10.3390/cells12050685.

13. Lingasamy P, Tobi A, Kurm K, Tammik O, Teesalu T. Targeting oncofetal fibronectin and neuropilin-1 in solid tumors with PL2 peptide. Sci Rep 15: 29369, 2025. doi: 10.1038/s41598-025-11299-x.

14. Celià-Terrassa T, Kang Y. How important is EMT for cancer metastasis? PLoS Biol 22: e3002487, 2024. doi: 10.1371/journal.pbio.3002487.

15. Freire-de-Lima L, Gelfenbeyn K, Ding Y, Mandel U, Clausen H, Handa K, Hakomori SI. Involvement of O-glycosylation defining oncofetal fibronectin in epithelial-mesenchymal transition process. Proceedings of the National Academy of Sciences 108: 17690–17695, 2011. doi: 10.1073/pnas.1115191108.

16. dos Reis JS, da Costa Santos MAR, da Costa KM, Freire-de-Lima CG, Morrot A, Previato JO, Previato LM, da Fonseca LM, Freire-de-Lima L. Increased expression of the pathological O-glycosylated form of oncofetal fibronectin in the multidrug resistance phenotype of cancer cells. Matrix Biology 118: 47–68, 2023. doi: 10.1016/j.matbio.2023.03.002.

17. David L, Mandel U, Clausen H, Sobrinho-Simões M. Immunohistochemical expression of oncofetal fibronectin in benign and malignant lesions of the stomach. Eur J Cancer 29: 2070– 2071, 1993. doi: 10.1016/0959-8049(93)90476-V.

18. Menzin AW, Loret de Mola JR, Bilker WB, Wheeler JE, Rubin SC, Feinberg RF. Identification of oncofetal fibronectin in patients with advanced epithelial ovarian cancer: detection in ascitic fluid and localization to primary sites and metastatic implants. Cancer 82: 152–8, 1998. doi: 10.1002/(sici)1097-0142(19980101)82:1<152::aid-cncr19>3.0.co;2-1.

19. Wunderlich H, Reichelt O, Zermann D-H, Schubert J, Berndt A, Kosmehl H. Fetal fibronectin: A new screening-marker for bladder cancer?.

20. Kaczmarek J, Castellani P, Nicolo G, Spina B, Allemanni G, Zardi L. Distribution of oncofetal fibronectin isoforms in normal, hyperplastic and neoplastic human breast tissues. Int J Cancer 59: 11–16, 1994. doi: 10.1002/ijc.2910590104.

21. Hirane N, Yokoi Y, Kobayashi A, Yamada M, Konno T, Kumeta H, Wakui H, Otaki M, Miura Y, Kakisaka T, Kamiyama T, Taketomi A, Nouso K, Kimura S, Nishimura S-I. Site-Specific O-Glycosylation in Oncofetal Fibronectin IIICS Domain Creates Cancer Stage-Specific Biomarkers. J Am Chem Soc 147: 12170–12184, 2025. doi: 10.1021/jacs.5c00333.

22. King-Peoples T, Keane JT, Lewis H-GS, Liu F, Brookens SK, Dagher OK, Parvathaneni K, Gardner BL, Ahmed A, Furth EE, Lal P, Posey Jr AD. CAR T Cells Targeting O-Glycosylated Fibronectin Exhibit Potent Cytolytic Activity and Combine with Tumoral Toll-Like Receptor Agonism to Overcome Tumor Resistance. Cancer Immunol Res 14: 1400–1415, 2026.

23. Hynes RO. The Extracellular Matrix: Not Just Pretty Fibrils. Science (1979) 326: 1216–1219, 2009. doi: 10.1126/science.1176009.

24. Saraswathibhatla A, Indana D, Chaudhuri O. Cell–extracellular matrix mechanotransduction in 3D. Nat Rev Mol Cell Biol 24: 495–516, 2023. doi: 10.1038/s41580-023-00583-1.

25. Kechagia JZ, Ivaska J, Roca-Cusachs P. Integrins as biomechanical sensors of the microenvironment. Nat Rev Mol Cell Biol 20: 457–473, 2019. doi: 10.1038/s41580-019-0134-2.

26. Ho KKY, Buschhaus JM, Zhang A, Cutter AC, Humphries BA, Luker GD. Substrate stiffness regulates triple-negative breast cancer signaling through CXCR4 receptor dynamics. Sci Rep 15: 29621, 2025. doi: 10.1038/s41598-025-14495-x.

27. Park H-J, Helfman DM. Up-regulated fibronectin in 3D culture facilitates spreading of triple negative breast cancer cells on 2D through integrin β-5 and Src. Sci Rep 9: 19950, 2019. doi: 10.1038/s41598-019-56276-3.

28. García-Rodríguez JL, Korsgaard U, Vissing SM, Paasch TP, Semenova M, Vendelbo SL, Jensby EF, Williams HL, Salachan PV, Brandt CB, Hanimann J, Lin L, Zlobec I, Sørensen KD, Kjems J, Hager H, Kristensen LS. Cancer-associated fibroblasts shape the formation of budding cancer cells at the invasive front of human colorectal cancer. Commun Biol 8: 1345, 2025. doi: 10.1038/s42003-025-08799-x.

29. Barbazan J, Pérez-González C, Gómez-González M, Dedenon M, Richon S, Latorre E, Serra M, Mariani P, Descroix S, Sens P, Trepat X, Vignjevic DM. Cancer-associated fibroblasts actively compress cancer cells and modulate mechanotransduction. Nat Commun 14: 6966, 2023. doi: 10.1038/s41467-023-42382-4.

30. Attieh Y, Clark AG, Grass C, Richon S, Pocard M, Mariani P, Elkhatib N, Betz T, Gurchenkov B, Vignjevic DM. Cancer-associated fibroblasts lead tumor invasion through integrin-β3– dependent fibronectin assembly. Journal of Cell Biology 216: 3509–3520, 2017. doi: 10.1083/jcb.201702033.

31. Erdogan B, Ao M, White LM, Means AL, Brewer BM, Yang L, Washington MK, Shi C, Franco OE, Weaver AM, Hayward SW, Li D, Webb DJ. Cancer-associated fibroblasts promote directional cancer cell migration by aligning fibronectin. Journal of Cell Biology 216: 3799–3816, 2017. doi: 10.1083/jcb.201704053.

32. Berger AJ, Renner CM, Hale I, Yang X, Ponik SM, Weisman PS, Masters KS, Kreeger PK. Scaffold stiffness influences breast cancer cell invasion via EGFR-linked Mena upregulation and matrix remodeling. Matrix Biology 85–86: 80–93, 2020. doi: 10.1016/j.matbio.2019.07.006.

33. Bordeleau F, Califano JP, Negrón Abril YL, Mason BN, LaValley DJ, Shin SJ, Weiss RS, Reinhart-King CA. Tissue stiffness regulates serine/arginine-rich protein-mediated splicing of the extra domain B-fibronectin isoform in tumors. Proceedings of the National Academy of Sciences 112: 8314–8319, 2015. doi: 10.1073/pnas.1505421112.

34. Efthymiou G, Radwanska A, Grapa AI, Beghelli-De la Forest Divonne S, Grall D, Schaub S, Hattab M, Pisano S, Poet M, Pisani DF, Counillon L, Descombes X, Blanc-Feraud L, Van Obberghen-Schilling E. Fibronectin Extra Domains tune cellular responses and confer topographically distinct features to fibril networks. J Cell Sci 134, 2021. doi: 10.1242/jcs.252957.

35. Liao YF, Gotwals PJ, Koteliansky VE, Sheppard D, Van Water L De. The EIIIA segment of fibronectin is a ligand for integrins α9β1 and α9β1 providing a novel mechanism for regulating cell adhesion by alternative splicing. Journal of Biological Chemistry 277: 14467–14474, 2002. doi: 10.1074/jbc.M201100200.

36. Sens C, Huck K, Pettera S, Uebel S, Wabnitz G, Moser M, Nakchbandi IA. Fibronectins containing extradomain A or B enhance osteoblast differentiation via distinct integrins. Journal of Biological Chemistry 292: 7745–7760, 2017. doi: 10.1074/jbc.M116.739987.

37. Rodrigues J, Heinrich MA, Teixeira LM, Prakash J. 3D In Vitro Model (R)evolution: Unveiling Tumor–Stroma Interactions. Trends Cancer 7: 249–264, 2021. doi: 10.1016/j.trecan.2020.10.009.

38. Miyauchi Y, Yasuchika K, Fukumitsu K, Ishii T, Ogiso S, Minami T, Kojima H, Yamaoka R, Katayama H, Kawai T, Yoshitoshi-Uebayashi EY, Kita S, Yasuda K, Sasaki N, Uemoto S. A novel three-dimensional culture system maintaining the physiological extracellular matrix of fibrotic model livers accelerates progression of hepatocellular carcinoma cells. Sci Rep 7, 2017. doi: 10.1038/s41598-017-09391-y.

39. Grey JFE, Campbell-Ritchie A, Everitt NM, Fezovich AJ, Wheatley SP. The use of decellularised animal tissue to study disseminating cancer cells. J Cell Sci 132, 2019. doi: 10.1242/jcs.219907.

40. Wishart AL, Conner SJ, Guarin JR, Fatherree JP, Peng Y, Mcginn RA, Crews R, Naber SP, Hunter M, Greenberg AS, Oudin MJ. Decellularized extracellular matrix scaffolds identify full-length collagen VI as a driver of breast cancer cell invasion in obesity and metastasis [Online]. Sci Adv 6, 2020. https://www.science.org.

41. Ferreira LP, Gaspar VM, Mano JF. Decellularized Extracellular Matrix for Bioengineering Physiomimetic 3D in Vitro Tumor Models. Trends Biotechnol. 38 Elsevier Ltd: 1397–1414, 2020.

42. Castelló-Cros R, Cukierman E. Stromagenesis During Tumorigenesis: Characterization of Tumor-associated Fibroblasts and Stroma-derived 3D Matrices [Online]. In: Extracellular Matrix Protocols, Methods in Molecular Biology, p. 275–305. www.springer.com/series/7651.

43. Wang K, Wu F, Seo BR, Fischbach C, Chen W, Hsu L, Gourdon D. Breast cancer cells alter the dynamics of stromal fibronectin-collagen interactions. Matrix Biology 60–61: 86–95, 2017. doi: 10.1016/j.matbio.2016.08.001.

44. Syed S, Karadaghy A, Zustiak S. Simple Polyacrylamide-based Multiwell Stiffness Assay for the Study of Stiffness-dependent Cell Responses. Journal of Visualized Experiments, 2015. doi: 10.3791/52643.

45. Tse JR, Engler AJ. Preparation of Hydrogel Substrates with Tunable Mechanical Properties. Curr Protoc Cell Biol 47, 2010. doi: 10.1002/0471143030.cb1016s47.

46. Young DA, Choi YS, Engler AJ, Christman KL. Stimulation of adipogenesis of adult adipose-derived stem cells using substrates that mimic the stiffness of adipose tissue. Biomaterials 34: 8581–8588, 2013. doi: 10.1016/j.biomaterials.2013.07.103.

47. Wang K, Andresen Eguiluz RC, Wu F, Seo BR, Fischbach C, Gourdon D. Stiffening and unfolding of early deposited-fibronectin increase proangiogenic factor secretion by breast cancer-associated stromal cells. Biomaterials 54: 63–71, 2015. doi: 10.1016/j.biomaterials.2015.03.019.

48. Franco-Barraza J, Raghavan KS, Luong T, Cukierman E. Engineering clinically-relevant human fibroblastic cell-derived extracellular matrices. In: Methods in Cell Biology, edited by Caballero D, Kundu SC, Luís Reis R. 2020, p. 109–160.

49. Malik R, Luong T, Cao X, Han B, Shah N, Franco-Barraza J, Han L, Shenoy VB, Lelkes PI, Cukierman E. Rigidity controls human desmoplastic matrix anisotropy to enable pancreatic cancer cell spread via extracellular signal-regulated kinase 2. Matrix Biology 81: 50–69, 2019. doi: 10.1016/j.matbio.2018.11.001.

50. Stringer C, Wang T, Michaelos M, Pachitariu M. Cellpose: a generalist algorithm for cellular segmentation. Nat Methods 18: 100–106, 2021. doi: 10.1038/s41592-020-01018-x.

51. Schindelin J, Arganda-Carreras I, Frise E, Kaynig V, Longair M, Pietzsch T, Preibisch S, Rueden C, Saalfeld S, Schmid B, Tinevez J-Y, White DJ, Hartenstein V, Eliceiri K, Tomancak P, Cardona A. Fiji: an open-source platform for biological-image analysis. Nat Methods 9: 676– 682, 2012. doi: 10.1038/nmeth.2019.

52. Papes F, Camargo AP, de Souza JS, Carvalho VMA, Szeto RA, LaMontagne E, Teixeira JR, Avansini SH, Sánchez-Sánchez SM, Nakahara TS, Santo CN, Wu W, Yao H, Araújo BMP, Velho PENF, Haddad GG, Muotri AR. Transcription Factor 4 loss-of-function is associated with deficits in progenitor proliferation and cortical neuron content. Nat Commun 13: 2387, 2022. doi: 10.1038/s41467-022-29942-w.

53. Liu Y, Keikhosravi A, Mehta GS, Drifka CR, Eliceiri KW. Methods for Quantifying Fibrillar Collagen Alignment. 2017, p. 429–451.

54. Maziveyi M, Dong S, Baranwal S, Alahari SK. Nischarin regulates focal adhesion and Invadopodia formation in breast cancer cells. Mol Cancer 17: 21, 2018. doi: 10.1186/s12943-018-0764-6.

55. Aditya V, Tambe V, Yue W. Development of a Novel Automated Workflow in Fiji ImageJ for Batch Analysis of Confocal Imaging Data to Quantify Protein Colocalization Using Manders Coefficient. Bio Protoc 15, 2025. doi: 10.21769/BioProtoc.5285.

56. Sahai E, Astsaturov I, Cukierman E, DeNardo DG, Egeblad M, Evans RM, Fearon D, Greten FR, Hingorani SR, Hunter T, Hynes RO, Jain RK, Janowitz T, Jorgensen C, Kimmelman AC, Kolonin MG, Maki RG, Powers RS, Puré E, Ramirez DC, Scherz-Shouval R, Sherman MH, Stewart S, Tlsty TD, Tuveson DA, Watt FM, Weaver V, Weeraratna AT, Werb Z. A framework for advancing our understanding of cancer-associated fibroblasts. Nat Rev Cancer 20: 174–186, 2020. doi: 10.1038/s41568-019-0238-1.

57. Singer II, Kawka DW, Kazazis DM, Clark RA. In vivo co-distribution of fibronectin and actin fibers in granulation tissue: immunofluorescence and electron microscope studies of the fibronexus at the myofibroblast surface. J Cell Biol 98: 2091–2106, 1984. doi: 10.1083/jcb.98.6.2091.

58. Bloom L, Ingham KC, Hynes RO. Fibronectin Regulates Assembly of Actin Filaments and Focal Contacts in Cultured Cells via the Heparin-binding Site in Repeat III _13_. Mol Biol Cell 10: 1521– 1536, 1999. doi: 10.1091/mbc.10.5.1521.

59. Miron-Mendoza M, Graham E, Manohar S, Petroll WM. Fibroblast-fibronectin patterning and network formation in 3D fibrin matrices. Matrix Biology 64: 69–80, 2017. doi: 10.1016/j.matbio.2017.06.001.

60. Simon T, Salhia B. Cancer-Associated Fibroblast Subpopulations With Diverse and Dynamic Roles in the Tumor Microenvironment. Molecular Cancer Research 20: 183–192, 2022. doi: 10.1158/1541-7786.MCR-21-0282.

61. Patel AK, Vipparthi K, Thatikonda V, Arun I, Bhattacharjee S, Sharan R, Arun P, Singh S. A subtype of cancer-associated fibroblasts with lower expression of alpha-smooth muscle actin suppresses stemness through BMP4 in oral carcinoma. Oncogenesis 7: 78, 2018. doi: 10.1038/s41389-018-0087-x.

62. Forsthuber A, Aschenbrenner B, Korosec A, Jacob T, Annusver K, Krajic N, Kholodniuk D, Frech S, Zhu S, Purkhauser K, Lipp K, Werner F, Nguyen V, Griss J, Bauer W, Soler Cardona A, Weber B, Weninger W, Gesslbauer B, Staud C, Nedomansky J, Radtke C, Wagner SN, Petzelbauer P, Kasper M, Lichtenberger BM. Cancer-associated fibroblast subtypes modulate the tumor-immune microenvironment and are associated with skin cancer malignancy. Nat Commun 15: 9678, 2024. doi: 10.1038/s41467-024-53908-9.

63. Schütz S, Solé-Boldo L, Lucena-Porcel C, Hoffmann J, Brobeil A, Lonsdorf AS, Rodríguez-Paredes M, Lyko F. Functionally distinct cancer-associated fibroblast subpopulations establish a tumor promoting environment in squamous cell carcinoma. Nat Commun 14: 5413, 2023. doi: 10.1038/s41467-023-41141-9.

64. Muchlińska A, Nagel A, Popęda M, Szade J, Niemira M, Zieliński J, Skokowski J, Bednarz-Knoll N, Żaczek AJ. Alpha-smooth muscle actin-positive cancer-associated fibroblasts secreting osteopontin promote growth of luminal breast cancer. Cell Mol Biol Lett 27: 45, 2022. doi: 10.1186/s11658-022-00351-7.

65. Chen J, Yang P, Xiao Y, Zhang Y, Liu J, Xie D, Cai M, Zhang X. Overexpression of α-sma-positive fibroblasts (CAFs) in Nasopharyngeal Carcinoma Predicts Poor Prognosis. J Cancer 8: 3897–3902, 2017. doi: 10.7150/jca.20324.

66. Talele NP, Fradette J, Davies JE, Kapus A, Hinz B. Expression of α-Smooth Muscle Actin Determines the Fate of Mesenchymal Stromal Cells. Stem Cell Reports 4: 1016–1030, 2015. doi: 10.1016/j.stemcr.2015.05.004.

67. Bieback K, Ha VA-T, Hecker A, Grassl M, Kinzebach S, Solz H, Sticht C, Klüter H, Bugert P. Altered Gene Expression in Human Adipose Stem Cells Cultured with Fetal Bovine Serum Compared to Human Supplements. Tissue Eng Part A 16: 3467–3484, 2010. doi: 10.1089/ten.tea.2009.0727.

68. Kirk T, Ahmed A, Rognoni E. Fibroblast Memory in Development, Homeostasis and Disease. Cells 10: 2840, 2021. doi: 10.3390/cells10112840.

69. Emon B, Joy MSH, Lalonde L, Ghrayeb A, Doha U, Ladehoff L, Brockstein R, Saengow C, Ewoldt RH, Saif MTA. Nuclear deformation regulates YAP dynamics in cancer associated fibroblasts. Acta Biomater 173: 93–108, 2024. doi: 10.1016/j.actbio.2023.11.015.

70. Calvo F, Ege N, Grande-Garcia A, Hooper S, Jenkins RP, Chaudhry SI, Harrington K, Williamson P, Moeendarbary E, Charras G, Sahai E. Mechanotransduction and YAP-dependent matrix remodelling is required for the generation and maintenance of cancer-associated fibroblasts. Nat Cell Biol 15: 637–646, 2013. doi: 10.1038/ncb2756.

71. Elosegui-Artola A, Andreu I, Beedle AEM, Lezamiz A, Uroz M, Kosmalska AJ, Oria R, Kechagia JZ, Rico-Lastres P, Le Roux A-L, Shanahan CM, Trepat X, Navajas D, Garcia-Manyes S, Roca-Cusachs P. Force Triggers YAP Nuclear Entry by Regulating Transport across Nuclear Pores. Cell 171: 1397–1410.e14, 2017. doi: 10.1016/j.cell.2017.10.008.

72. Driscoll TP, Cosgrove BD, Heo S-J, Shurden ZE, Mauck RL. Cytoskeletal to Nuclear Strain Transfer Regulates YAP Signaling in Mesenchymal Stem Cells. Biophys J 108: 2783–2793, 2015. doi: 10.1016/j.bpj.2015.05.010.

73. Belhabib I, Zaghdoudi S, Lac C, Bousquet C, Jean C. Extracellular Matrices and Cancer-Associated Fibroblasts: Targets for Cancer Diagnosis and Therapy? Cancers (Basel) 13: 3466, 2021. doi: 10.3390/cancers13143466.

74. Busch S, Landberg G. CAF-specific markers: role of the TGFβ pathway. Oncoscience 2: 835– 836, 2015. doi: 10.18632/oncoscience.209.

75. Ringuette Goulet C, Bernard G, Tremblay S, Chabaud S, Bolduc S, Pouliot F. Exosomes Induce Fibroblast Differentiation into Cancer-Associated Fibroblasts through TGFβ Signaling. Molecular Cancer Research 16: 1196–1204, 2018. doi: 10.1158/1541-7786.MCR-17-0784.

76. Ding L, Li Z, Yue J, Qiu L, Tian Z, Zhang H. Cancer-associated fibroblasts as a critical driver in tumor metastasis: The mechanisms and future perspectives. iScience 29: 114788, 2026. doi: 10.1016/j.isci.2026.114788.

77. Ulmer J, Geiger B, Spatz JP. Force-induced fibronectin fibrillogenesis in vitro. Soft Matter 4: 1998, 2008. doi: 10.1039/b808020h.

78. Erickson HP. Reversible unfolding of fibronectin type III and immunoglobulin domains provides the structural basis for stretch and elasticity of titin and fibronectin. Proceedings of the National Academy of Sciences 91: 10114–10118, 1994. doi: 10.1073/pnas.91.21.10114.

79. Oakes PW, Beckham Y, Stricker J, Gardel ML. Tension is required but not sufficient for focal adhesion maturation without a stress fiber template. Journal of Cell Biology 196: 363–374, 2012. doi: 10.1083/jcb.201107042.

80. Boscher C, Dennis JW, Nabi IR. Glycosylation, galectins and cellular signaling. Curr Opin Cell Biol 23: 383–392, 2011. doi: 10.1016/j.ceb.2011.05.001.

81. Giannini R, Faviana P, Cavinato T, Elisei R, Pacini F, Berti P, Fontanini G, Ugolini C, Camacci T, De Ieso K, Miccoli P, Pinchera A, Basolo F. Galectin-3 and Oncofetal-Fibronectin Expression in Thyroid Neoplasia as Assessed by Reverse Transcription-Polymerase Chain Reaction and Immunochemistry in Cytologic and Pathologic Specimens. Thyroid 13: 765–770, 2003. doi: 10.1089/105072503768499662.

82. Reticker-Flynn NE, Malta DFB, Winslow MM, Lamar JM, Xu MJ, Underhill GH, Hynes RO, Jacks TE, Bhatia SN. A combinatorial extracellular matrix platform identifies cell-extracellular matrix interactions that correlate with metastasis. Nat Commun 3: 1122, 2012. doi: 10.1038/ncomms2128.

83. Reticker-Flynn NE, Bhatia SN. Aberrant Glycosylation Promotes Lung Cancer Metastasis through Adhesion to Galectins in the Metastatic Niche. Cancer Discov 5: 168–181, 2015. doi: 10.1158/2159-8290.CD-13-0760.

84. Lagana A, Goetz JG, Cheung P, Raz A, Dennis JW, Nabi IR. Galectin Binding to Mgat5-Modified N-Glycans Regulates Fibronectin Matrix Remodeling in Tumor Cells. Mol Cell Biol 26: 3181–3193, 2006. doi: 10.1128/MCB.26.8.3181-3193.2006.

